# Microbial food web dynamics in tropical waters of the bluefin tuna spawning region off northwestern Australia

**DOI:** 10.1101/2025.08.24.672009

**Authors:** Michael R. Landry, Michael R. Stukel, Natalia Yingling, Karen E. Selph, Sven A. Kranz, Christian K. Fender, Rasmus Swalethorp, Ria I. Bhabu

## Abstract

Whereas recruitment success for many fisheries depends on coincident timing of larvae with abundance peaks of their prey, less can be more in the tropical/subtropical spawning areas of bluefin tunas if lower but steady food resources are offset by reduced larval vulnerability to pelagic predators. To understand larval habitat characteristics for Southern Bluefin Tuna (SBT), we quantified microbial community carbon flows based on growth and grazing rates from depth profiles of dilution incubations and carbon biomass assessments from microscopy and flow cytometry (FCM) during their peak spawning off NW Australia (Indian Ocean) in February 2022. Two Chl*a*-based estimates of phytoplankton production gave differing offsets due to cycling or mixotrophy, exceeding ^14^C net community production on average (677 ± 98 versus 447 ± 43 mg C m^−2^ d^−1^). Productivity was higher than in the Gulf of Mexico spawning area for Atlantic Bluefin Tuna but less than similar studies of oceanic upwelling regions. Microzooplankton grazing averaged 482 ± 63 mg C m^−2^ d^−1^ (71 ± 13% of production). Two measurement variables for *Prochlorococcus* gave average production and grazing rates of 282 ± 36 and 248 ± 32 mg C m^−2^ d^−1^ (86 ± 6% grazed). *Prochlorococcus* comprised almost half of production and grazing fluxes in the upper (0-25 m) euphotic zone where SBT larvae reside. *Prochlorococcus* declined and eukaryotic phytoplankton and heterotrophic bacteria increased in relative importance in the lower euphotic zone. These results describe relatively classic open-ocean oligotrophic conditions as the food web base for nutritional flows to SBT larvae.

## 1. Introduction

The small area of the eastern Indian Ocean between northwestern Australia and Indonesia and downstream of the Indonesian Throughflow is a region of global significance as the only low-latitude connection between major oceans, the pathway through which excess heat and nutrients from the western Pacific are transferred to the Indian Ocean (Lee et al., 2015; Desbruyéres et al., 2017), and the only known spawning region for Southern Bluefin Tuna (SBT) (Matsuura et al., 1997; Farley and Davis, 1998). It is also, however, a very sparsely studied area, with few biogeochemical or ecological measurements to relate to other tropical/subtropical systems, to evaluate impacts of the Pacific exchange, to understand how the production environment functions in general and as critical habitat for larval tuna, or to assess potential vulnerabilities of the ecosystem and larval habitat to future climate change. In January-February 2022, the BLOOFINZ (Bluefin Larvae in Oligotrophic Ocean Foodwebs, Investigations of Nutrients to Zooplankton) Program conducted field sampling and experimental studies in this region during the peak SBT spawning season to advance the knowledge base for evaluating such issues.

As part of the BLOOFINZ-IO (Indian Ocean) expedition, we investigated microbial food web dynamics – the growth and mortality interactions of protists and bacteria that generate and consume most of the ocean’s productivity (Azam et al., 1983; Rivkin and Legendre, 2001; Calbet and Landry, 2004). We used depth profiles of dilution incubations to determine instantaneous rates of growth and grazing for phytoplankton and bacterial components measured by pigments and flow cytometry. Carbon-based estimates of food web fluxes combined these measured rates with microscopic and flow cytometric biomass assessments and provided a basis for comparison to other tropical/subtropical systems that have been studied similarly (e.g., Landry et al., 2011, 2016, 2022a,b).

The relative magnitudes and distributions of C flows within microbial food webs are relevant to the growth and survival of larval tuna in two opposing ways. Rich systems with more direct and efficient nutritional pathways to mesozooplankton (i.e., proportionally greater C flows from larger primary producers, like diatoms) can enhance larval prey resources that promote faster growth. At the same time, higher mesozooplankton concentrations can support other predators, small schooling pelagics in particular, that can greatly diminish the survival potential of larvae (Bakun and Broad, 2003; Bakun, 2013). The tradeoffs in this growth-survival conundrum suggest a delicate balance in larval habitat quality, leaning toward warm oligotrophic systems, where food web fluxes support sufficient prey for adequate to rapid growth while minimizing vulnerability to predators (Shropshire et al., 2022). As the first step to understanding the lower level food web dynamics supporting the SBT spawning region, we consider the relative contributions of *Prochlorococcus*, eukaryotic protists, microzooplankton grazers and heterotrophic bacteria to production and C flows, we validate community production relative to contemporaneous ^14^C net primary production, we compare to other systems, and we distinguish food web scenarios for the upper euphotic zone (EZ) where larval tuna reside versus the extended production environment of the lower EZ that supplements nutrition of their zooplankton prey.

## 2. Materials and methods

### 2.1. Sampling and experimental set-up

Four experimental studies of phytoplankton growth and microzooplankton grazing were conducted on *R/V Roger Revelle* cruise RR2201 in January-March 2022 in waters overlying the Argo Abyssal Plain, a deep oceanic basin off NW Australia (hereafter, Argo Basin). Each major experiment, called a “cycle”, involved a repeated daily sequence of water-column sampling and process rate measurements following a satellite-tracked free-drifting array (Landry et al., 2009) consisting of a surface float, a 3-m drogue centered at 15 m, a coated-wire with stainless-steel attachment rings for *in situ* bottle incubations, and a separately attached smaller float with iridium transmission (10-min position frequency) and nighttime strobe light. Cycle 1 (C1) was conducted from 3-7 February in W-SW flowing water in the SW corner of Argo Basin (Fig. 1). C2 was conducted from 9-13 February in E-SE flowing water overlying the south basin slope. C3 was conducted from 15-18 February in northward flowing water to the east of C1, beginning at the location of a drifter released at the end of C1 (Fig. 1). C4 was conducted from 20-24 February in an area of weak S-SW flow in the SE corner of Argo Basin.

**Fig. 1.**
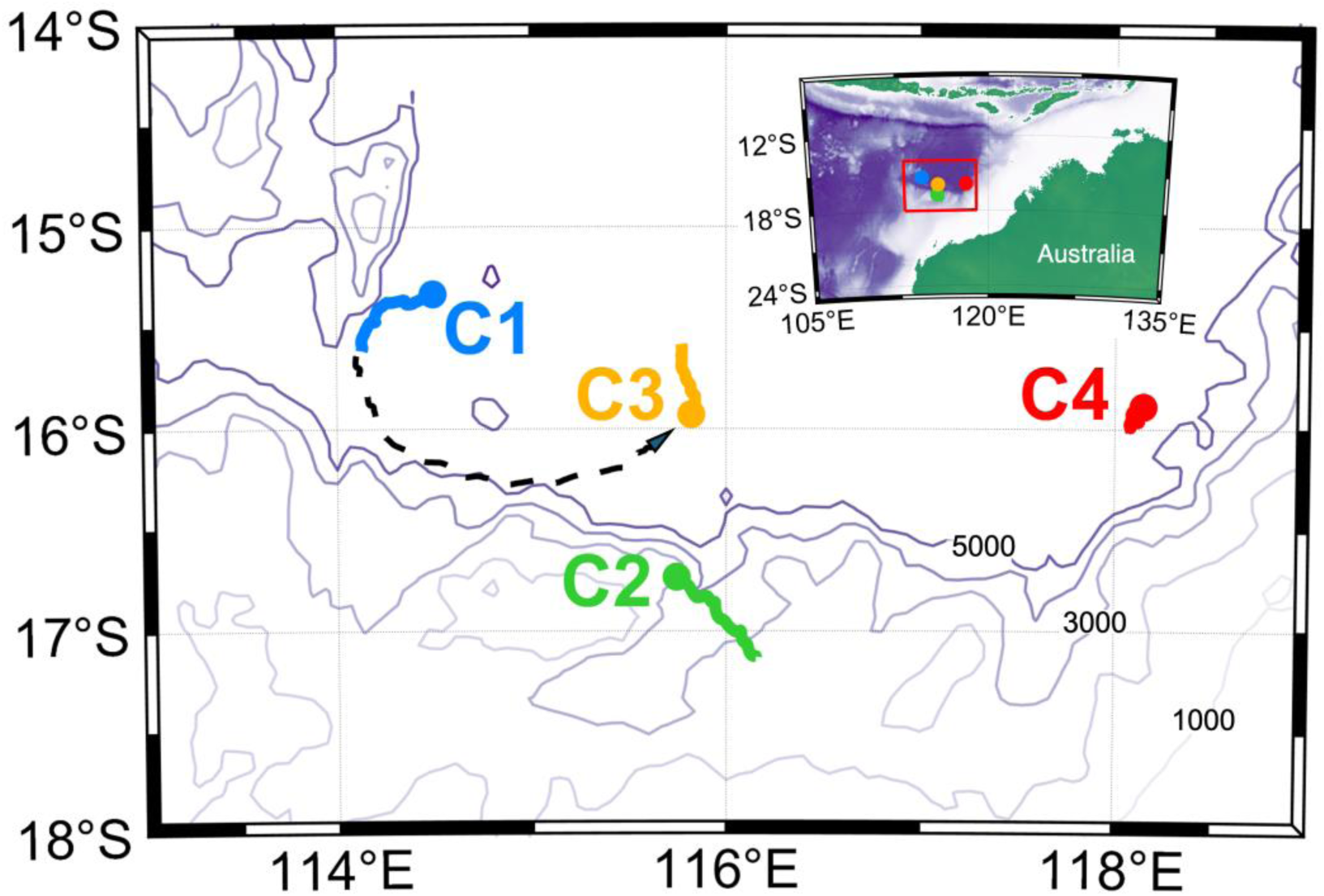
Map of the study region off NW Australia showing starting locations (colored dots) and drifter trajectories for four Cycle experiments (C1 to C4) conducted in the southern Argo Basin. Dashed line shows path of a drogued drifter that was deployed at the end of C1 and recovered for the start of C3. Bathymetric contours are plotted at 1000-m depth intervals.

For each cycle experiment, we collected seawater daily from Niskin bottles on early-morning CTD hydrocasts (∼02:00 local time) at 6 depths, ranging from 5 m to the deep chlorophyll maximum (DCM) in the lower euphotic zone. Initial samples for pigment concentrations, flow cytometry (FCM) and microscopy were taken directly from the Niskin bottles via silicone tubing. For each depth sampled, we also prepared a two-treatment dilution experiment (Landry et al., 2008, 2011), with one polycarbonate bottle (2.7 L) containing unfiltered seawater (100%) and the second (diluted) bottle consisting of ∼33% whole seawater with filtered water from the same depth. Seawater was filtered directly from the Niskin bottles using a peristaltic pump, silicone tubing and an in-line 0.2 µm Suporcap (Pall Acro) filter capsule that had previously been acid washed (3.7% trace-metal grade HCl; Milli-Q and seawater rinses). Dilution bottles were first given a measured volume of filtered water and then gently filled to the top with unscreened water from the Niskin bottles to minimize damage to fragile protists (Gifford, 1988; Lessard and Murrell, 1998). Consistent with previous studies with the two-treatment method in open-ocean systems (Landry et al., 2008, 2009, 2011, 2016), nutrients were not added to the incubation bottles. Lessard and Murrell (1998) demonstrated that added nutrients were not needed for reasonable rate estimates and linearity in dilution experiments conducted in the oligotrophic Sargasso Sea, whereas adding nutrients resulted in erratic results and depressed grazing. After preparation, each bottle was subsampled for flow cytometric (FCM) analysis (1 mL) for initial concentrations and volumetric dilution factors. In addition to the dilution experiment bottles, 280-mL polycarbonate bottles (triplicate “light”, one “dark”) were filled at each depth and spiked with H^14^CO ^−^ to measure net community productivity (NCP; for additional details see Kranz et al., this issue).

All bottles were secured in coarse net bags clipped to 1-m separated upper and lower attachment rings on the drifter line and incubated *in situ* for 24 h at the depth of collection. For the first deployment of each cycle, the entire array with bottle bags attached was laid out on deck before being quickly lowered by hand. For daily experiments, we set up a new experiment with water collected in close proximity (∼100 m) to the drifter position before recovering the drifter. The drifter was then hand recovered, the previous day’s experiments removed, the new experiments attached, and the array redeployed – a process that took 10-15 min while the ship maintained position. All recovery and deployments were carried out before sunrise. Upon recovery, all bottles were subsampled for flow cytometry and fluorometric Chl*a* (FlChl*a* hereafter), and the remaining volume (2.1 L) was filtered for pigment analyses by high-performance liquid chromatography (HPLC).

### 2.2. Environmental measurements

Environmental measurements were mainly averaged or integrated from CTD instrument readings at 1-m resolution for temperature, salinity, σ_T_ density, light (photosynthetically available radiation, PAR) and *in situ* fluorescence. Mixed-layer depth (MLD) is defined as the depth at which σ_T_ density first exceeded the 0-5 m average by 0.01 kg m^−3^. The deep chlorophyll maximum (DCM) is the depth of maximum profile fluorescence. Euphotic zone (EZ) depth is the depth of penetration of 1% surface irradiance, computed from mean light extinction coefficients in the upper 5-100 m. Nitrate concentrations are from CTD water samples (50-mL prewashed HDPE bottles) filtered through the 0.2-µm Suporcap filter capsule during experimental setup, frozen at −20°C and subsequently analyzed on a Seal Analytical continuous-flow AutoAnalyzer 3 by the SIO Oceanographic Data Facility (full data presented in Kranz et al., this issue).

### 2.3. Pigment analyses

Initial and final samples (250 mL) for FlChl*a* were filtered onto GF/F filters and extracted with 90% acetone in a −20°C freezer for 24 h. Extracted samples were warmed to room temperature in the dark and analyzed on a Turner Designs model 10 fluorometer calibrated against a pure Chl*a* standard (Strickland and Parsons, 1972).

Samples (2.3 L for initials; 2.1 L for finals) for HPLC analyses of chlorophyll and carotenoid pigments were concentrated onto Whatman GF/F filters under low vacuum pressure, immediately frozen in liquid nitrogen and stored at −80°C. The samples were extracted for 2 h in 100% methanol, disrupted by sonication, clarified by GF/F filtration and analyzed (Agilent Technologies 1200 Series) at the analytical facility of the Institut de la Mer de Villefranche (CNRS-France) according to procedures described in Ras et al. (2008).

### 2.4. Flow cytometry and microscopical analyses

Flow cytometric (FCM) and epifluorescence (EPI) microscopical analyses of the microbial community were done as described by Yingling et al. (this issue). FCM samples (1 mL) were preserved with 0.5% paraformaldehyde (v/v) and stored in the dark at 4°C until shipboard analysis, typically within several hours of collection. The samples were first stained with Hoechst 34580 (1 µg mL^−1^) then enumerated at a flow rate of 50 µL min^−1^ with a Beckman-Coulter CytoFLEX S cytometer with 4 lasers (Selph, 2021). Side scatter, forward angle light scatter (FALS) and fluorescence signals were measured using laser excitation (EX)/emission (EM) filters of EX375/EM450 ± 45 for Hoechst-stained DNA, EX488/EM690 ± 50 for chlorophyll, and EX561/EM585 ± 42 for phycoerythrin. Listmode files (FCS 3.0) were analyzed with FlowJo software (v.10.6.1) for abundances of *Prochlorococcus* (PRO), *Synechococcus* (SYN), photosynthetic eukaryotes (PEUK) and heterotrophic bacteria (HBAC), as well as their population-average fluorescence and scatter signals normalized to fluorescent bead standards. Bacterial cell abundances were converted to carbon biomass using factors of 5 fg C cell^−1^ for HBAC (Gunderson et al., 2002; Landry et al., 2023) and 32 and 101 fg C cell^−1^ for PRO and SYN, respectively (Garrison et al., 2000; Brown et al., 2008).

Seawater samples (450 mL) for analysis by epifluorescence microscopy (EPI) were preserved with sequential additions of alkaline Lugol’s solution, borax-buffered formaldehyde and sodium thiosulfate (method of Sherr and Sherr, 1993 as modified by Taylor et al., 2015), and stained with 1 mL of proflavin (0.33% w/v) and 1 mL of DAPI (0.01 mg mL^−1^) prior to filtering. Subsamples of 50 mL (SV = small volume slides) were filtered onto 25-mm, black, 0.8-µm pore polycarbonate filters to enumerate cells with equivalent spherical diameters (ESD) of 12 µm or less. The remaining 400 mL (LV = large volume slides) was filtered onto 25-mm, black, 8.0-µm pore polycarbonate filters to enumerate cells with >12-µm ESD. Each filter was mounted onto a glass slide using Type DF immersion oil and a No. 2 cover slip.

The slides were imaged with an Olympus Microscope DP72 Camera on an Olympus BX51 fluorescence microscope (Yinging et al., this issue). Thirty nanoplankton images (60x objective lens) and 20 microplankton images (20x) were captured using filter sets for proflavine (482 nm excitation, 536 nm emission – cell protein), chlorophyll autofluorescence (450-490 nm excitation, 660-680 nm emission) and DAPI (350 nm excitation, 465 nm emission – cell DNA). Images were individually identified as phototrophic or heterotrophic based on chlorophyll presence and sized with ImageJ software. Cell biovolumes (BV) were calculated from the formula for a prolate sphere, BV = 0.5236×L×W^2^, where L is measured cell length (µm) on the polar axis and W is measured width on the equatorial axis (Taylor et al., 2016). Carbon (C; pg cell^−1^) biomass was computed from the equations: C = 0.216×BV^0.939^ for non-diatoms, and C = 0.288×BV^0.811^ for diatoms (Menden-Deuer and Lessard, 2000). The abundance difference between FCM-enumerated PEUK cells and chlorophyll-containing cells on the SV slides and PEUK cells were assumed to be the eukaryotic picophytoplankton (<2-µm cells) missed by microscopy (Taylor and Landry, 2018) and were added (320 fg C cell^−1^) to the total sample biomass.

Seawater samples (250 mL) were also preserved with 5% acid Lugol’s solution for microscopical analyses of ciliates by the Utermöhl method (Lund et al., 1958). For each daily profile, samples from the upper three sampling depths were volumetrically combined, following the trapezoidal rule, to produce one sample representing the upper EZ, and the lower three samples were combined to produce one sample for the lower EZ. The combined samples were measured for total volume in a volumetric cylinder, settled for 24 h, concentrated by suctioning off the upper water and ultimately settled and analyzed in the Utermöhl chamber. Recognizable ciliates and diatoms were enumerated for the full settled samples with a Zeiss AxioVert 200 M inverted microscope at 200X magnification and brightfield illumination. The first 50-60 cells of each group were imaged and sized for mean BV estimates using FIJI software (Schindelin et al., 2012). For ciliates, we calculated carbon biomass as pg C = 0.19×BV (Putt and Stoecker, 1989). C biomasses of diatoms are from the equation of Menden-Deuer and Lessard (2000).

### 2.5. Microbial growth and grazing rates

We determined rate profiles for phytoplankton growth (µ, d^−1^) and microzooplankton grazing (m, d^−1^) from each pair of dilution experiment bottles and FCM or pigment-associated population according to the following equations:

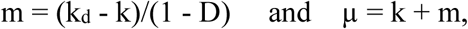

where k_d_ and k are the measured net rates of change between initial and final concentrations in the diluted and undiluted treatments, respectively, and D is the portion of unfiltered water in the dilution treatment, measured as the mean ratio of PRO cell abundances in initial diluted and undilute samples from FCM analyses (Landry et al., 2008; Selph et al., 2011). Rate estimates assume comparable growth rates in dilution treatments and proportional grazing relative to dilution, consistent with the expected close coupling of production, grazing and nutrient remineralization in the microbial communities of oligotrophic systems. Rate estimates for PRO and HBAC cell abundances are from FCM analyses. Pigment-based rates are from FlChl*a* or individual HPLC chlorophylls or carotenoids, with growth rates corrected for changes in cellular chlorophyll [ln(RF_final_/RF_init_)], where RF is mean bead-normalized red fluorescence of PRO or PEUK cells from FCM analyses (Landry et al., 2003). We used initial and final RF of PRO cells to correct growth rates from divinyl Chl*a* (DVChl*a*), and initial and final RF of PEUK cells to correct growth rates from monovinyl Chl*a* (MVChl*a*) and carotenoid-based rates for individual groups of eukaryotic phytoplankton. Total community growth rates from FlChl*a* or total HPLC Chl*a* (TChl*a* hereafter) were corrected based on the proportional contributions of PRO and PEUK cells to total RF.

Some experiments at individual depths were compromised by a bottle loss during incubation or sample loss during the filtration process. All of the 3^rd^ depth samples for HPLC samples also showed highly negative net rates in the diluted bottle for all pigment variables and all experiments, suggesting a systematic filtration leak for the large bottle samples at one position on the filtration rack. These experiments were excluded, leaving 67 experiments for rate comparisons among the different variables measured. Unless otherwise noted, we present experimental uncertainties as ± standard errors of mean estimates (± SEM), averaging by depth stratum and treating each cycle day as an independent experiment.

Carbon-based estimates of phytoplankton community production (PROD) and microzooplankton grazing (GRAZ) were calculated from growth (µ) and grazing (m) rates from dilution experiments and the following equations (Landry *et al*., 2000):

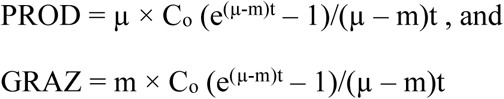

where C_o_ is initial autotrophic biomass (mg C m^−3^) and t = time (1 day). PROD and GRAZ estimates for the euphotic zone were determined by integrating rate estimates from the surface to the deepest incubation depth according to the trapezoidal rule.

## 3. Results

### 3.1. Environmental conditions

Environmental conditions for the cycle experiments can be summarized as warm, strongly stratified and oligotrophic (Table 1). Cycle 1 (C1) followed an early cruise storm and had the deepest mixed later depth (30 m) and slightly higher mean nitrate concentration in the upper 40 m, though still lower than 0.03 µM on average. Temporal interpretation of the water mass tracked from C1 to C3 experiments suggests that surface waters warmed, depths of the mixed layer, DCM and euphotic zone shallowed, and 0-80 m FlChl*a* increased over the two week period following storm mixing (Table 1). Warming of the upper EZ by ∼1.4°C is also evident as a temporal trend among all cycles, though mean temperature of the upper 100 m shows no change. Overall, despite some modest temporal or spatial variability, depth structure (DCM, EZ, nitracline) and mean environmental conditions remained relatively similar over the four experiments.

**Table 1.**
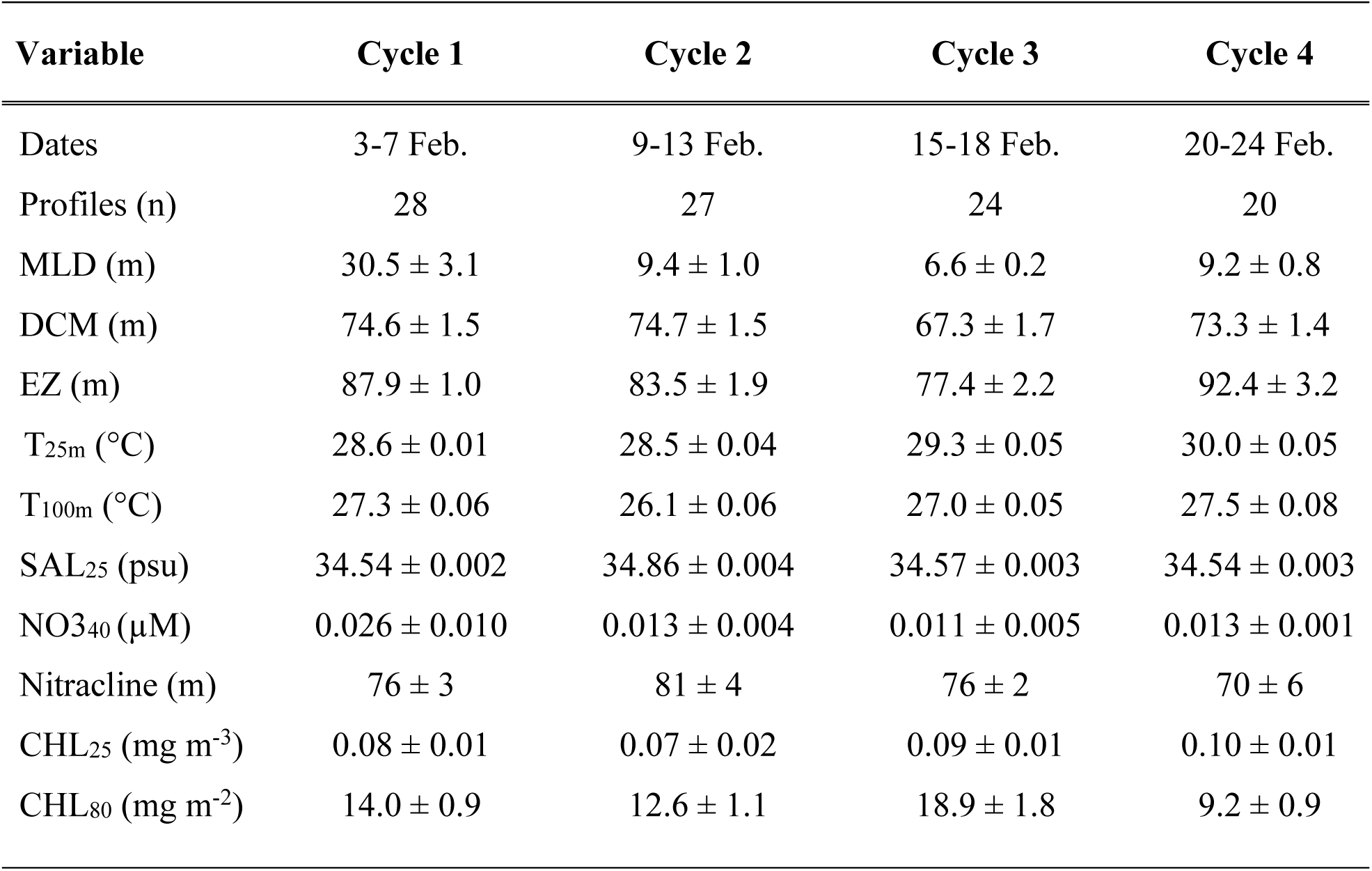
Environmental conditions for cycle experiments in the Argo Basin, NW Australia in February 2022. n = number of CTD profiles averaged. MLD (m) = Mixed Layer Depth, defined as the depth at which seawater density is 0.1 kg m^−3^ greater than mean upper 5 m. DCM (m) = depth of the Deep Chlorophyll Maximum; EZ (m) = depth of the euphotic zone (1% surface PAR); T_25_ (°C) = mean temperature of the mixed layer; T_100m_ (°C) = mean temperature of the upper 100 m; SAL_25_ (psu) = mean salinity of the mixed layer; NO3_40_ (µM) = mean nitrate concentration in the upper 40 m; and Nitracline (m) = mean depth of 1.0 µM nitrate concentration. CHL_25_ (mg m^−3^) and CHL_80_ (mg m^−2^) are average fluorometrically measured Chl*a* for the upper 25 m and integrated to 80 m, the mean depth of experimental incubations). Uncertainties are standard errors of mean values.

### 3.2. Pigment and FCM relationships

In general, the pigment and FCM variables for growth and grazing rate assessments showed strong relationships in this study region. Paired FlChl*a* and TChl*a* measurements gave a 1.0 slope (p<10^−90^, 95% confidence limits = 0.97, 1.09; Fig. 2A). Red fluorescences of the respective FCM populations were also well correlated with the summed pigment variables of TChl*a+b* (R=0.94, p<10^−114^; Fig. 2B), DVChl*a+b* (R=0.93; p<10^−103^; Fig. 2C) and MVChl*a+b* (R=0.94, p<10^−111^; Fig. 2D). Whereas RF for eukaryote cells was equally correlated with MVChl*a* as with MVChl*a+b*, using DVChl*a+b* instead of DVChl*a* as the plot variable notably improved the RF relationship for PRO cells because the deep EZ population of PRO had an order of magnitude higher ratio of DVChl*b* to DVChl*a* than the shallow population (0.68 versus 0.08).

**Fig. 2.**
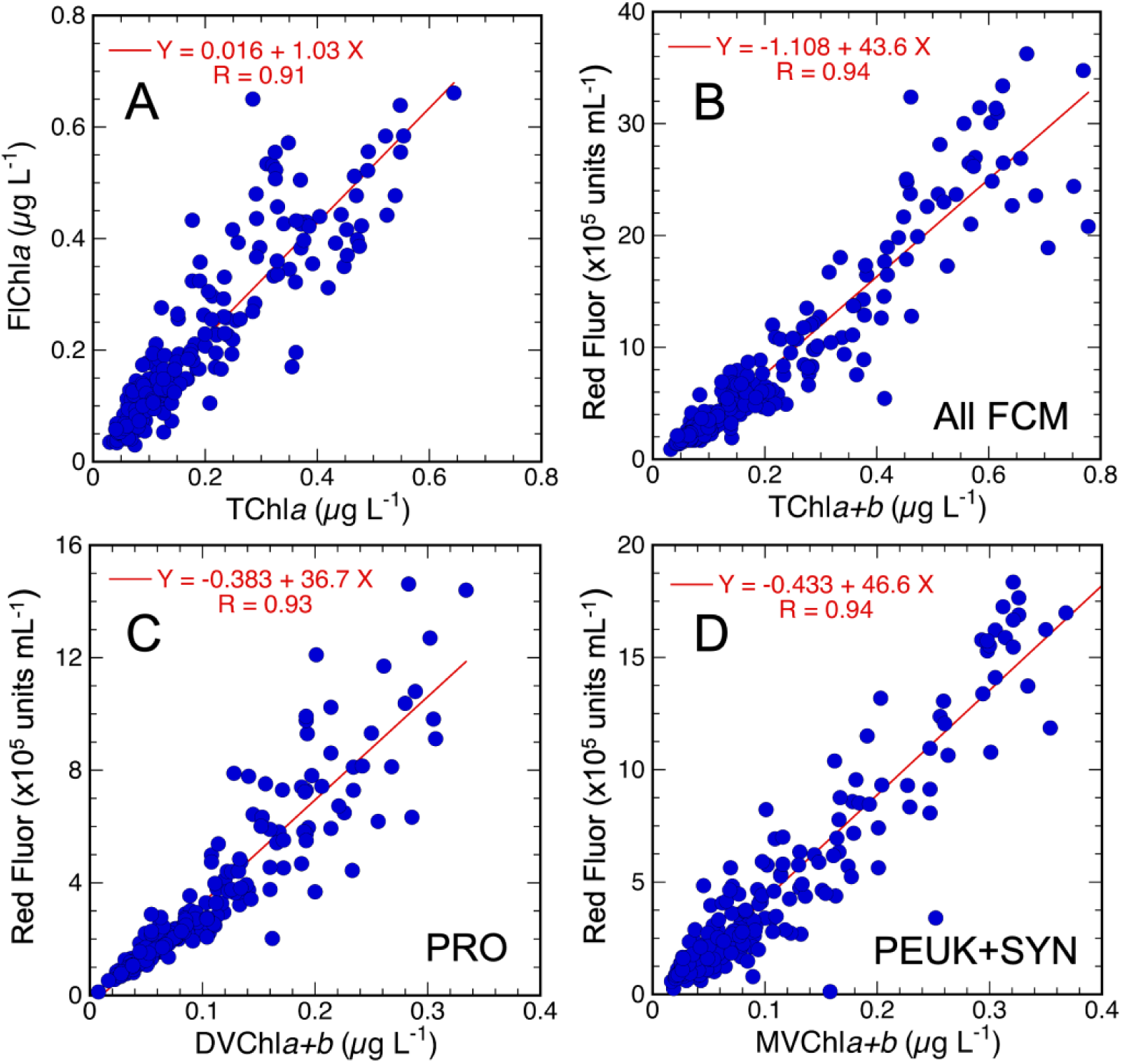
Relationships among measured pigment and flow cytometic (FCM) variables used to determine growth and grazing rates. Panel A: fluorometric estimates of Chl*a* (FlChl*a*) versus total Chla (TChl*a*) from HPLC. Panel B: total red fluorescence (Red Fluor, bead-calibrated units mL^−1^) from all FCM populations (*Prochlorococcus* – PRO, *Synechococcus* – SYN, and photosynthetic eukaryotes – PEUK) versus Total Chl*a* + Chl*b* from HPLC (TChl*a+b*). Panel C: total PRO Red Fluor mL^−1^ versus divinyl Chl*a* + Chl*b* from HPLC (DVChl*a+b*). Panel D: total PEUK+SYN Red Fluor mL^−1^ versus monovinyl Chl*a* + Chl*b* from HPLC (MVChl*a+b*). Data include all initial and final (post-incubation) samples with measurements by both methods.

### 3.3. Chlorophyll-based growth and grazing rates

Figure 3 compares mean cycle depth profiles of concentration, growth rate and grazing mortality as assessed from measurements of FlChl*a* and TChl*a* in the same incubation bottles. Consistent with the overall good relationship between these measured variables (Fig. 2A), the concentration profiles show similar features of low concentrations (<0.1 µg L^−1^) in the upper 20 m increasing to values of ∼0.3 to 0.45 µg L^−1^ in the deep EZ. For C1, however, the deep EZ estimates of FlChl*a* averaged 0.08 µg L^−1^ higher than for TChl*a* (p=0.014, paired t-test, two sided).

**Fig. 3.**
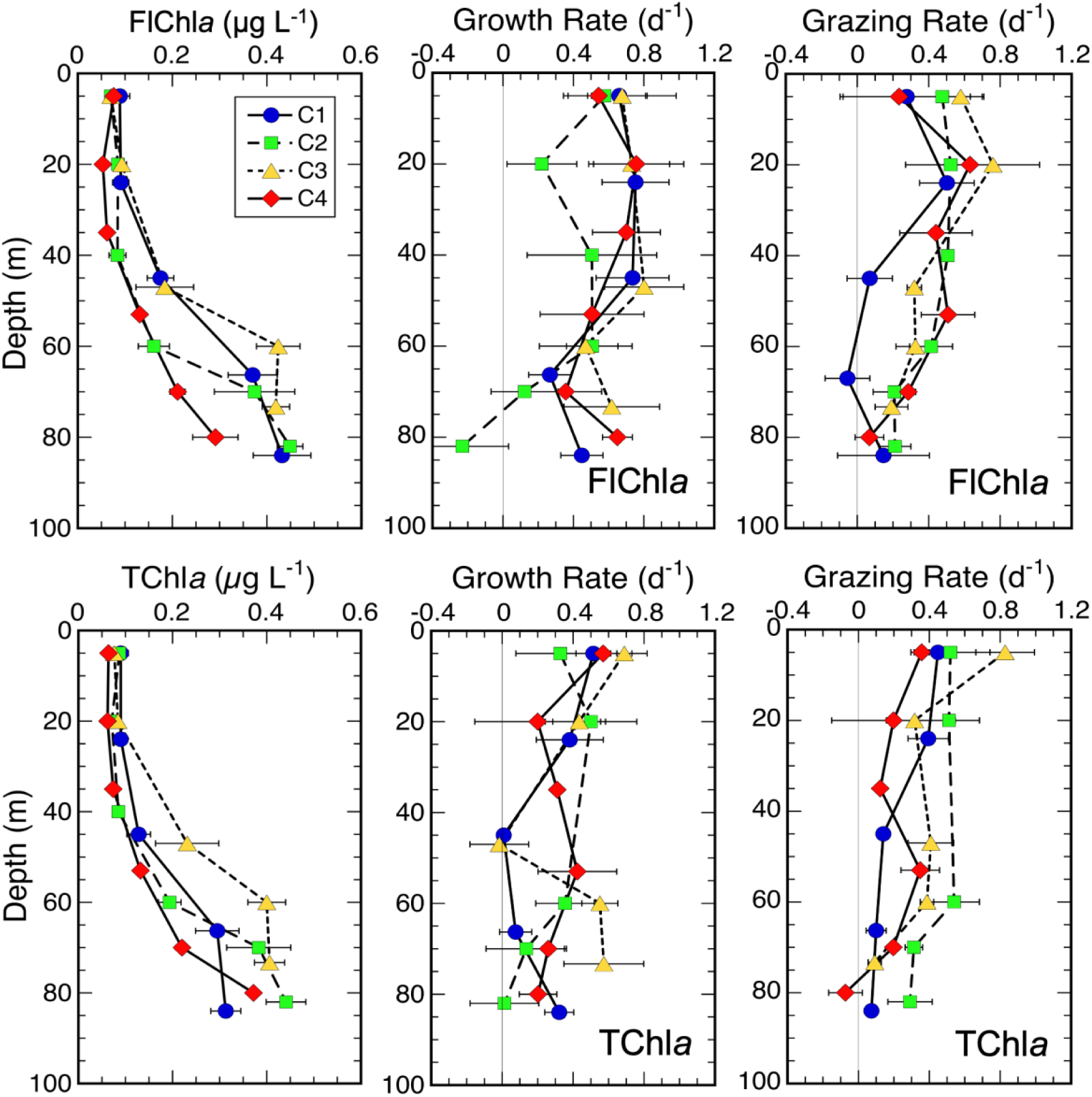
Mean depth profiles of Chl*a* concentrations, phytoplankton growth rates (d^−1^) and microzooplankton grazing mortality (d^−1^) from dilution experiments conducted during four experimental cycles (C1-C4). Top row: estimates from fluorometric measurements of Chl*a*. Bottom row: estimates from HPLC measurements of TChl*a*. Uncertainties are standard errors of mean values (SEM).

Compared to the overall similarities in concentration profiles, the mean cycle rate profiles for the two chlorophyll *a* methods are quite variable in magnitudes and shapes (Fig. 3). Cycle-averaged growth rates for the upper EZ (<25 m) ranged from 0.40 to 0.76 d^−1^ for FlChl*a* and from 0.35 to 0.56 d^−1^ for TChl*a*, with proportionately lower growth rates in the deep EZ (0.13-0.56 d^−1^ for FlChla, 0.15-0.37 d^−1^ for TChla) (Table 2). For grazing mortality, cycle-average rates ranged from 0.38 to 0.67 d^−1^ for FlChl*a* and from 0.26 to 0.57 d^−1^ for TChl*a* in the upper EZ and from 0.03 to 0.28 and 0.08 to 0.39 d^−1^, respectively, in the deep EZ (Fig. 3, Table 2). Averaged over all cycles, FlChl*a* growth rates were 34% higher than TChl*a* in the upper EZ (0.63 ± 0.07 versus 0.47 ± 0.06 d^−1^; p=0.10) and 35% higher in the lower EZ (0.36 ± 0.07 versus 0.24 ± 0.06 d^−1^, p=0.067), though neither of these differences were statistically significant (paired t-test, two sided). Overall rates for grazing mortality agreed closely between FlChl*a* and TChl*a* in both the upper (0.51 ± 0.07 versus 0.46 ± 0.06 d^−1^; p=0.64) and lower EZ (0.21 ± 0.06 versus 0.22 ± 0.04 d^−1^; p=0.57).

**Table 2.** Mean cycle rate estimates of growth (µ, d^−1^) and grazing mortality (m, d^−1^) from dilution incubations in the upper (<25 m) and lower (55-90 m) euphotic zone of the Argo Basin. Measured variables are fluorometric Chl*a* (FlChl*a*); total (TChl*a*), divinyl (DVChl*a*) and monovinyl (MVChl*a*), hexfucoxanthin (HEX), fucoxanthin (FUCO) and butfucoxanthin (BUT) from HPLC analyzed pigment samples; and flow cytometric cell counts of *Prochlorococcus* (PRO) and heterotrophic bacteria (HBAC). Uncertainties are standard errors of mean values.

| Variable | Cycle 1 |  | Cycle 2 |  | Cycle 3 |  | Cycle 4 |  |
| --- | --- | --- | --- | --- | --- | --- | --- | --- |
| | $\mu$ ($d^{-1}$) | $m$ ($d^{-1}$) | $\mu$ ($d^{-1}$) | $m$ ($d^{-1}$) | $\mu$ ($d^{-1}$) | $m$ ($d^{-1}$) | $\mu$ ($d^{-1}$) | $m$ ($d^{-1}$) |
| **Upper EZ (<25 m)** |  |  |  |  |  |  |  |  |
| FlChla | 0.76±0.09 | 0.38±0.17 | 0.40±0.16 | 0.50±0.16 | 0.70±0.17 | 0.67±0.14 | 0.67±0.16 | 0.47±0.14 |
| TChla | 0.53±0.05 | 0.44±0.06 | 0.43±0.17 | 0.51±0.12 | 0.56±0.09 | 0.57±0.15 | 0.35±0.23 | 0.26±0.20 |
| DVChla | 0.42±0.11 | 0.39±0.18 | 0.64±0.16 | 0.68±0.15 | 0.75±0.12 | 0.76±0.18 | 0.35±0.30 | 0.29±0.30 |
| MVChla | 0.68±0.13 | 0.38±0.07 | 0.19±0.22 | 0.31±0.11 | 0.40±0.18 | 0.36±0.14 | 0.28±0.20 | 0.22±0.10 |
| HEX | 0.66±0.20 | 0.56±0.24 | 0.39±0.31 | 0.37±0.22 | 0.75±0.19 | 0.48±0.19 | 0.33±0.26 | 0.11±0.10 |
| FUCO | 0.71±0.13 | 0.33±0.12 | 0.30±0.23 | 0.15±0.13 | 0.52±0.20 | 0.23±0.13 | 0.17±0.26 | 0.03±0.06 |
| BUT | 0.58±0.21 | 0.51±0.25 | 0.12±0.24 | 0.22±0.13 | 0.26±0.22 | 0.14±0.16 | -0.09±0.16 | -0.12±0.09 |
| PRO | 0.20±0.24 | 0.02±0.25 | 0.39±0.10 | 0.22±0.11 | 0.54±0.08 | 0.40±0.07 | 0.39±0.14 | 0.11±0.13 |
| HBAC | 0.25±0.07 | 0.22±0.08 | 0.33±0.06 | 0.23±0.03 | 0.17±0.05 | 0.14±0.06 | 0.03±0.03 | 0.01±0.04 |
| **Lower EZ (>55 m)** |  |  |  |  |  |  |  |  |
| FlChla | 0.32±0.08 | 0.03±0.11 | 0.13±0.15 | 0.28±0.07 | 0.56±0.12 | 0.28±0.05 | 0.52±0.11 | 0.27±0.09 |
| TChla | 0.15±0.06 | 0.08±0.02 | 0.17±0.11 | 0.39±0.07 | 0.37±0.14 | 0.30±0.05 | 0.30±0.08 | 0.15±0.08 |
| DVChla | 0.10±0.05 | 0.07±0.04 | 0.45±0.11 | 0.53±0.09 | 0.62±0.15 | 0.44±0.08 | 0.45±0.06 | 0.23±0.10 |
| MVChla | 0.12±0.07 | 0.09±0.03 | -0.05±0.13 | 0.26±0.07 | 0.22±0.16 | 0.22±0.07 | 0.21±0.11 | 0.09±0.09 |
| HEX | 0.07±0.07 | 0.10±0.03 | 0.03±0.11 | 0.25±0.10 | 0.15±0.14 | 0.13±0.14 | 0.17±0.15 | 0.01±0.12 |
| FUCO | 0.23±0.06 | 0.10±0.03 | 0.25±0.13 | 0.40±0.09 | 0.47±0.23 | 0.22±0.07 | 0.41±0.10 | 0.14±0.09 |
| BUT | 0.19±0.12 | 0.15±0.04 | 0.14±0.13 | 0.39±0.08 | 0.33±0.28 | 0.23±0.06 | 0.29±0.11 | 0.13±0.07 |
| PRO | 0.07±0.02 | 0.16±0.03 | 0.24±0.03 | 0.30±0.04 | 0.21±0.08 | 0.29±0.06 | 0.16±0.04 | 0.24±0.06 |
| HBAC | 0.34±0.07 | 0.24±0.07 | 0.48±0.11 | 0.40±0.12 | 0.17±0.03 | 0.12±0.04 | 0.13±0.03 | 0.09±0.03 |

The divinyl and monovinyl components of TChl*a* had relatively similar concentrations in the upper EZ but notably higher proportions of MVChl*a* in the lower EZ, indicating a greater contribution of eukaryotic phytoplankton (SYN abundances were low and contributed little to cell RF) relative to PRO in the DCM (Fig. 4). Mean-cycle growth rates varied from 0.35 to 0.75 d^−1^ for DVChl*a* and from 0.19 to 0.68 d^−1^ for MVChl*a* in the upper EZ, and declined to 0.10-0.62 d^−1^ and −0.05-0.22 d^−1^, respectively, in the deep EZ (Table 2). Grazing averages ranged from 0.29 to 0.76 d^−1^ for DVChl*a* and from 0.22 to 0.38 d^−1^ for MVChl*a* in the upper EZ and from 0.07 to 0.53 d^−1^ and 0.09 to 0.26 d^−1^, respectively, in the lower EZ (Fig. 3, Table 2). For all cycles, DVChl*a* rates in the upper EZ were 46% higher for growth (0.56 ± 0.08 versus 0.38 ± 0.09 d^−1^; p=0.21) and 64% higher for grazing (0.54 ± 0.09 versus 0.33 ± 0.05 d^−1^; p=0.023) than MVChl*a* rates, though only the grazing differences were significant. DVChl*a* estimates were significantly higher than MVChl*a* rates in the lower EZ for both growth (0.42 ± 0.05 versus 0.15 ± 0.06 d^−1^; p<10^−4^) and grazing (0.30 ± 0.06 versus 0.16 ± 0.04 d^−1^; p=0.0014).

**Fig. 4.**
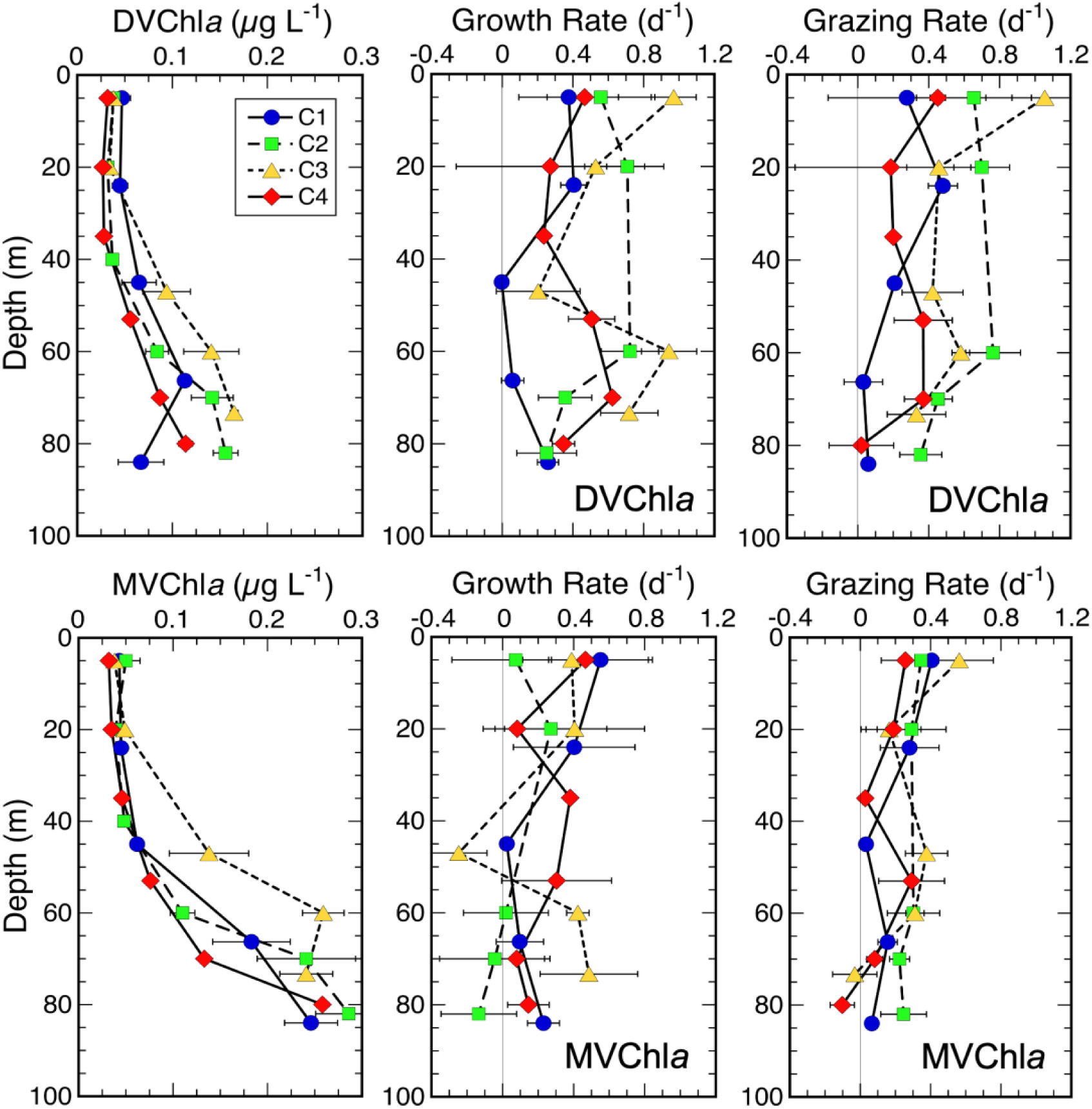
Mean depth profiles of Chl*a* concentrations, phytoplankton growth rates (d^−1^) and microzooplankton grazing mortality (d^−1^) from dilution experiments during experimental cycles (C1-C4). Top row: estimates from measurements of divinyl Chl*a*. Bottom row: estimates from measurements of monovinyl Chl*a*. Uncertainties are standard errors of mean values (SEM).

### 3.4. Rates estimates from group-specific carotenoids

Rate estimates for carotenoid pigments display high variability, including substantial excursions into negative values, and are best considered in terms of relative averages for the groups they represent (Fig. 5). HEX, representing prymnesiophytes and averaging 56% of total concentration of carotenoid pigments, had the highest mean growth and grazing estimates for the upper EZ (0.52 ± 0.11 and 0.39 ± 0.10 d^−1^, respectively). The other mean upper-EZ growth and grazing estimates are 0.48 ± 0.09 and 0.19 ± 0.06 d^−1^ for FUCO (diatoms) and 0.21 ± 0.09 and 0.20 ± 0.09 d^−1^ for BUT (pelagophytes). For the lower EZ, growth rates for HEX were notably reduced while grazing estimates were similar for all groups (0.12 ± 0.06 and 0.25 ± 0.12 d^−1^ for HEX; 0.37 ± 0.06 and 0.20 ± 0.06 d^−1^ for FUCO; 0.31 ± 0.07 and 0.25 ± 0.04 d^−1^ for BUT). FUCO had the highest positive growth-grazing difference (net growth) in both the upper and lower EZ.

**Fig. 5.**
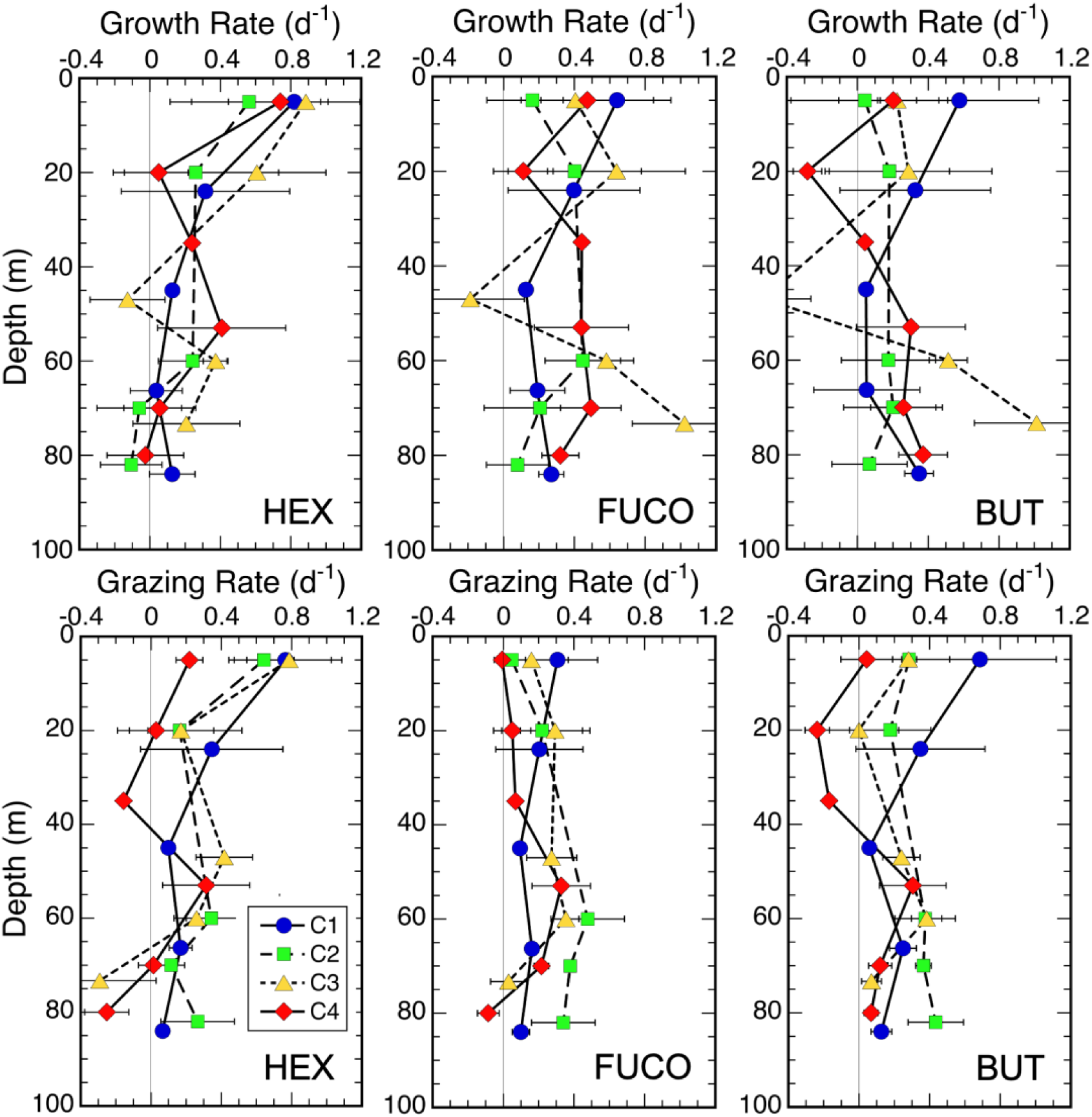
Mean depth profiles of growth rates (d^−1^) and microzooplankton grazing mortality (d^−1^) for phytoplankton group-specific carotenoid pigments during experimental cycles (C1-C4). HEX = 19’-hexanoyloxyfucoxanthin (prymnesiophytes); FUCO = fucoxanthin (diatoms); BUT = 19’-butanoyloxyfucoxanthin (pelagophytes). Uncertainties are standard errors of mean values (SEM).

### 3.5. Bacterial rate estimates from flow cytometry

Abundance profiles for PRO showed relatively consistent densities of 2-3×10^5^ cells mL^−1^ with modest mid-EZ peaks for C2-C4 (Fig. 6). HBAC densities were relatively uniform with depth (∼5×10^5^ cells mL^−1^) for C1 and C2 but had deep EZ maxima of ∼9×10^5^ cells mL^−1^ for C3 and C4. For PRO, cycle-averaged growth rates for the upper EZ ranged from 0.20 to 0.54 d^−1^ and were always higher than their respective cycle-averaged grazing estimates (range 0.02 to 0.40 d^−1^; Table 2). In the lower EZ, however, mean rates of grazing loss (range 0.16-0.30 d^−1^) exceeded cell growth rates of PRO (range 0.07-0.24 d^−1^; Table 2) for all cycles. For HBAC, mean lower-EZ growth (range 0.13 to 0.48 d^−1^) exceeded upper-EZ growth rate (range 0.03 to 0.33 d^−1^) for all cycles, and grazing mortality rates for both the upper (range 0.01-0.23 d^−1^) and lower EZ (range 0.09-0.40 d^−1^) left small net positive growth differences with respect to their corresponding cycle growth-rate averages.

**Fig. 6.**
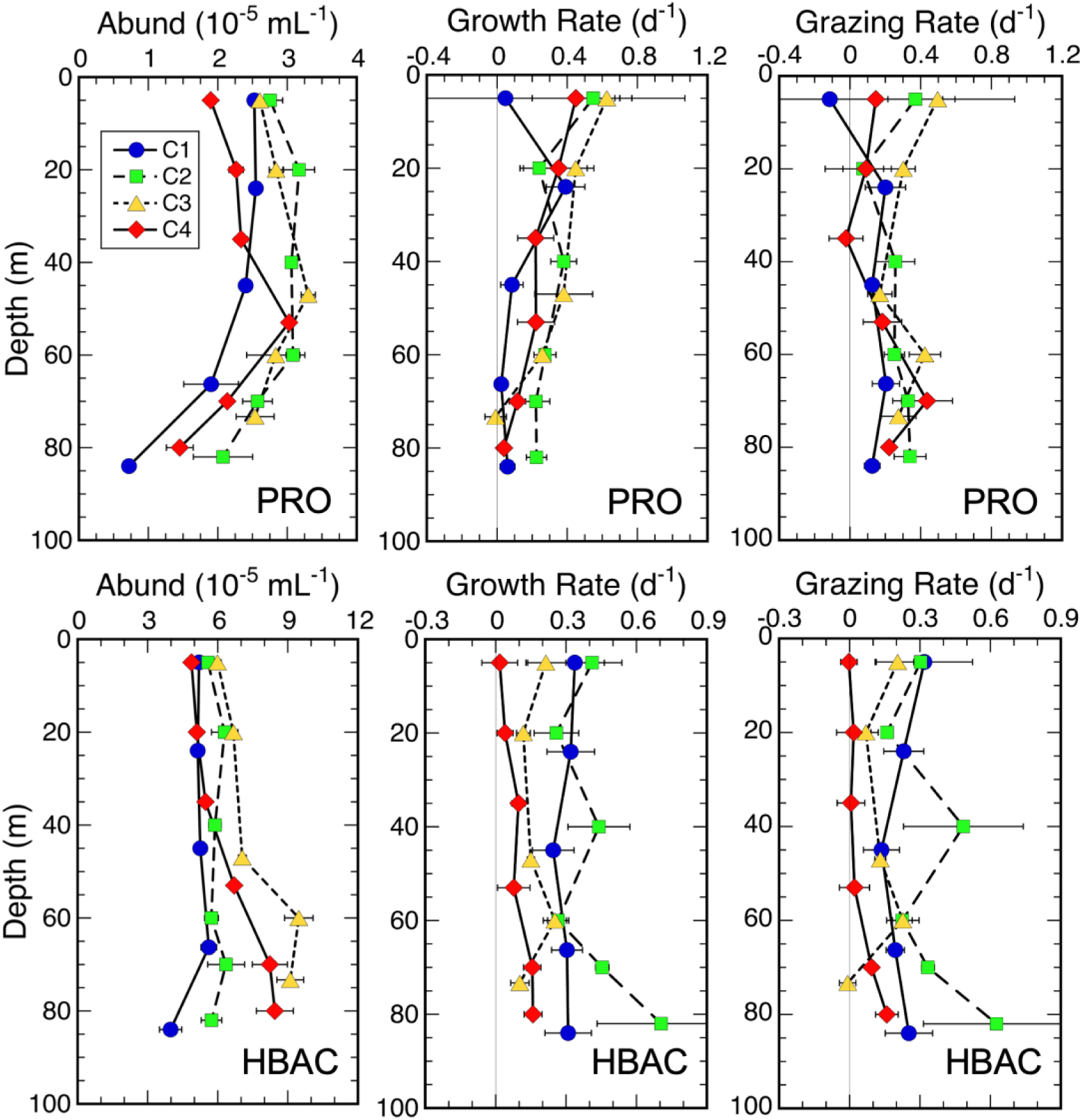
Mean depth profiles of cell abundances, growth rates (d^−1^) and microzooplankton grazing mortality (d^−1^) for *Prochlorococcus* (PRO) and heterotrophic bacteria (HBAC) during experimental cycles (C1-C4). Uncertainties are standard errors of mean values (SEM).

### 3.6. EZ-integrated production and grazing

Carbon-based rates of production (PROD) and grazing (GRAZ) in Table 3 were calculated from instantaneous µ and m rate estimates and the portions of phytoplankton community C corresponding to those rates (total phytoplankton C for FlChl*a* and TChl*a* rates; PRO, HBAC and PEUK biomass for DVChla, PRO, HBAC and MVChl*a* rates). Carbon rates were first calculated for each incubation depth then integrated for daily profiles and averaged for each cycle. We treated negative rates as zeros in this process, resulting in slightly higher (3%) mean estimates of depth-integrated carbon than the same data with negative rates, but eliminating issues with negative grazing:growth ratios for calculated estimates of percent production consumed.

**Table 3.** Carbon-based estimates of integrated production (PROD) and grazing (GRAZ) to the depths of experimental incubations. ^14^C-NCP = Net Community Production from ^14^C *in situ* incubations on the drifter array (Kranz et al., this volume). FlChl*a* and TChl*a* are total phytoplankton community rates from fluorometric and HPLC measures of chlorophyll *a*, respectively. MVChl*a* and DVChl*a* are partial community rates for *Synechococcus* + eukaryotes (cells with monovinyl Chl*a*) and *Prochloroccus* (cells with divinyl Chl*a*), respectively. PRO_fcm and HBAC_fcm are rates from flow cytometric cell counts of *Prochlorococcus* and heterotrophic bacteria, respectively. Uncertainties are standard errors of mean values.

| Variable | Cycle 1 | Cycle 2 | Cycle 3 | Cycle 4 | All |
| --- | --- | --- | --- | --- | --- |
| **PROD (mg C m<sup>-2</sup> d<sup>-1</sup>)** |  |  |  |  |  |
| $^{14}\text{C}$-NCP | 262 ± 66 | 528 ± 46 | 540 ± 52 | 517 ± 23 | 447 ± 43 |
| FlChla | 969 ± 193 | 574 ± 165 | 982 ± 249 | 815 ± 114 | 772 ± 108 |
| TChla | 413 ± 130 | 580 ± 272 | 631 ± 386 | 491 ± 126 | 527 ± 118 |
| MVChla | 385 ± 149 | 182 ± 129 | 363 ± 286 | 125 ± 73 | 258 ± 81 |
| DVChla | 167 ± 28 | 499 ± 140 | 401 ± 118 | 377 ± 84 | 340 ± 58 |
| PRO | 123 ± 40 | 261 ± 35 | 259 ± 34 | 201 ± 77 | 199 ± 25 |
| HBAC | 71 ± 9 | 94 ± 7 | 42 ± 2 | 30 ± 5 | 63 ± 8 |
| **GRAZ (mg C m<sup>-2</sup> d<sup>-1</sup>)** |  |  |  |  |  |
| FlChla | 393 ± 30 | 618 ± 193 | 551 ± 111 | 463 ± 234 | 492 ± 67 |
| TChla | 333 ± 36 | 660 ± 165 | 402 ± 29 | 367 ± 132 | 455 ± 68 |
| MVChla | 180 ± 53 | 178 ± 59 | 120 ± 24 | 96 ± 80 | 142 ± 26 |
| DVChla | 174 ± 21 | 517 ± 118 | 341 ± 26 | 308 ± 27 | 320 ± 51 |
| PRO | 105 ± 31 | 207 ± 25 | 196 ± 14 | 109 ± 1 | 155 ± 17 |
| HBAC | 53 ± 7 | 77 ± 9 | 28 ± 10 | 19 ± 3 | 47 ± 7 |

For the total phytoplankton community, production estimates from FlChl*a* ranged from 574 to 982 mg C m^−2^ d^−1^ for individual cycles (mean 772 ± 108 mg C m^−2^ d^−1^) and were generally higher than those from TChl*a* (range 413 to 631 mg C m^−2^ d^−1^, mean 527 ± 118 mg C m^−2^ d^−1^; p=0.008, paired t-test, 2-sided), except for C2 (Table 3). TChl*a* estimates were closest to the measured rates from *in situ* ^14^C-NCP incubations for each cycle and overall (range 262 to 540 mg C m^−2^ d^−1^, mean 447 ± 43 mg C m^−2^ d^−1^). Compared to production, integrated grazing estimates from FlChl*a* and TChl*a* were similar in cycle ranges (393-618 versus 333-660 mg C m^−2^ d^−1^) and overall averages (492 ± 67 versus 455 ± 68 mg C m^−2^ d^−1^; p=0.23) (Table 3), giving lower cycle-mean estimates of % production grazed by microzooplankton for FlChl*a* than TChl*a* (65 ± 15 versus 83 ± 11 %).

For the monovinyl and divinyl components of TChl*a*, C-based rate estimates from eukaryote-dominated MVChl*a* were lower than those from PRO-associated DVChl*a*, but only significantly so for grazing (142 ± 26 versus 320 ± 51 mg C m^−2^ d^−1^; p=0.003) and not production (258 ± 81 versus 340 ± 58 mg C m^−2^ d^−1^; p=0.40) (Table 3). Accordingly, a lower percentage of eukaryote production was consumed by microzooplankton compared to consumption of PRO production (63 ± 15 versus 94 ± 6 %). However, the production and grazing estimates for PRO based on FCM cell counts were lower than DVChl*a*-based rates for all cycles and for the experiment averages (production: 209 ± 25 versus 356 ± 61 mg C m^−2^ d^−1^, p=0.03; grazing: 158 ± 18 versus 339 ± 52 mg C m^−2^ d^−1^, p=0.001), and the FCM rates gave a lower percentage of PRO production consumed (74 ± 7 % versus 94 ± 6 %).

Compared to phytoplankton, C-based rate estimates for HBAC were relatively low (Table 3). Production and grazing rates averaged 63 ± 8 mg C m^−2^ d^−1^ (range: 30 to 94 mg C m^−2^ d^−1^) and 47 ± 7 mg C m^−2^ d^−1^ (range: 19 to 77 mg C m^−2^ d^−1^), respectively. From the cycle means, microzooplankton consumed 79 ± 5 % of HBAC production, on average.

## 4. Discussion

Aside from modest near-surface warming and intensifying stratification, four experimental studies of phytoplankton growth and microzooplankton in the southern Argo Basin were conducted under relatively similar environmental conditions, with low nutrient concentrations extending to at least 40 m and deep euphotic zones, deep nitraclines and prominent DCMs (Table 1). For these relatively classic open-ocean oligotrophic conditions, no phytoplankton group substantially exceeded cycle-averaged growth rates of one cell doubling d^−1^ (0.69 d^−1^) in the upper EZ, and the overall means were less (highest for FlChl*a* = 0.63 ± 0.07 d^−1^) (Table 2). Growth rate estimates typically declined by about 2 fold in the deep EZ (depths corresponding to the upper shoulder to maximum of the DCM), and a large proportion of production for all measured variables was consumed by micrograzers in both the upper and lower EZ. In the Discussion sections below, we refine rate estimates further with constraints from related measured variables, compare them to results from other relevant systems and apply them to microbial community C flux syntheses for the upper and lower EZ.

### 4.1. Rate issues and interpretations

Two prominent issues arise as observed rate discrepancies between alternate measures of the same phytoplankton group (FlChl*a* and TChl*a* as alternate indices of total phytoplankton biomass, and DVChl*a* and FCM cell counts as alternate measures of PRO). For the former, C production estimates from FlChl*a* were significantly higher than those from TChl*a* but grazing estimates were more similar, giving rise to two different scenarios of the percentage of phytoplankton production consumed by micro-grazers (65 versus 83%). Since we used the same cell RF corrections and C biomass data for these two variables, the differences come from the instantaneous rate estimates of µ and m. For the two PRO-associated variables, rate estimates from DVChl*a* were substantially higher than those from FCM cell counts, and two different scenarios of microzooplankton grazing impact were also evident as higher percentage grazing from DVChl*a* (96 versus 74%). For both measured variables, we used the same C biomass estimates for PRO in the production and grazing calculations. While these primary µ and m estimates come from different approaches, they are linked in part by using measured RF change in PRO cells FCM analyses to correct DVChl*a* rates.

The FlChl*a*-TChl*a* rate discrepancy is informed by comparing integrated C-based estimates of phytoplankton production from those variables to ^14^C-NCP results from the same incubation conditions (Table 4). Dilution estimates of production are expected to be higher than ^14^C-NCP rates, particularly for warm-water high-turnover environments, due to PO^14^C losses to community cycling. From JGOFS studies in the Arabian Sea, Dickson et al. (2001) found that ^14^C uptake rates from 24-h incubations averaged 21% lower than those for 12-h (end of daytime) incubations. From a grazing model for the same program, Laws et al. (2000) concluded that grazing alone, at a rate of 0.69 d^−1^ (equivalent to removing growth of one cell doubling d^−1^), would reduce ^14^C counts by 15% over 24 h. Consistent with these conclusions (accounting for both daytime grazing and cycling, and bulk nighttime loss), EZ-integrated production estimates from TChl*a* in warm waters of the equatorial Pacific were 29% higher than 24-h ^14^C-NCP (Landry et al., 2011). Production estimates based on the growth rates of plastidic (nominally “phytoplankton”) cells can also exceed ^14^C production estimates if some of those cells have a mixed nutritional mode (mixotrophy) in which a some portion of growth comes from ingestion of prey that are weakly ^14^C labelled (Mitra et al., 2014). Intuitively, TChl*a* would seem to be the more reliable metric for dilution net-change calculations because it is a high-precision measure of absolute Chl*a* concentration compared to FlChl*a*, which is calculated from two (initial and acidified) sample readings that are sensitive to presence of other pigments, like Chl*b*. In the present data, however, TChl*a* production estimates were closer to ^14^C-NCP (15% difference) than they realistically should be to account for community cycling losses, while FlChl*a* estimates at the same time were substantially higher (possibly indicative of mixotrophy). Given these differences and their implications, it is reasonable to retain features of both variables by averaging the integrated rates from each experiment as a blended product. The blended mean production of the phytoplankton community is 677 ± 98 mg C m^−2^ d^−1^. The blended mean grazing is 482 ± 63 mg C m^−2^ d^−1^, giving an average of 71 ± 13 % of phytoplankton production grazed by microzooplankton.

Given the above rationale for averaging alternate estimates of production and grazing for the full phytoplankton community, a similar approach also seems reasonable for the alternate estimates of PRO production and grazing. Both of the measurement variables, DVChl*a* and FCM cell counts, are from high-precision methods that have shown similar rate estimates in previous direct comparisons (e.g., Selph et al., 2011; Landry et al., 2011), but there is no independent metric to distinguish between them when they differ. Mean upper-EZ growth rates from PRO cell counts in the Argo Basin (0.39 d^−1^) are on the low side of estimates from the same method in other tropical/subtropical systems, more typically in the range of 0.5-0.7 d^−1^ (Landry et al., 2011, 2016, 2022a,b for equatorial Pacific, Costa Rica Dome, Gulf of Mexico and eastern Indian Ocean, respectively). Growth rates from cell counts might be underestimated if some growth over the incubation period is hidden as increased mean cell biomass or larger cells in the diluted treatment relative to the natural control. Using forward angle light scatter (FALS) as an index of cell size, FCM analyses provides some evidence of both effects, with mean-FALS rates of increase of 0.10 ± 0.01 d^−1^ in diluted and 0.04 ± 0.01 in undiluted treatments. If cell C content increases in proportion to FALS^0.55^ (DuRand and Olson, 1996), these cell biomass changes increase mean PRO µ by 0.07 d^−1^ and PRO m by 0.05 d^−1^. While the DVChl*a* estimates of PRO µ have a pigment correction that can also be questioned, they provide a coherent connection to pigment-based production and grazing estimates for the full community and the eukaryotic (MVChl*a*) component and are constrained by almost equally high rate estimates of grazing (PRO m) that are not affected by the pigment correction. The mean integrated values of PRO production and grazing from blended FCM and DVChl*a* rates are 282 ± 36 and 248 ± 32 mg C m^−2^ d^−1^, respectively, giving an overall estimate of 86 ± 6 % of PRO production grazed.

### 4.2. Comparisons to other subtropical and tropical ecosystems

With the Argo region being the only known global spawning region for southern bluefin tuna (SBT), it is useful to understand how lower trophic level processes there relate to other tropical/subtropical systems that have been studied similarly. In a previous investigation of the Gulf of Mexico spawning region for Atlantic bluefin tuna (Landry et al., 2022a), cycle-averaged estimates of FlChl*a*- and TChl*a*-based phytoplankton production (333-465 and 234-455 mg C m^−2^ d^−1^, respectively) were lower than same-variable estimates from Argo Basin (Table 3), but FCM-based estimates of PRO production (145-204 mg C m^−2^ d^−1^) fell in the same range as Argo values. Our blended estimates of PRO and total production give a mean contribution of PRO to total production similar to that in the oceanic GoM spawning region (42 versus 41%). In these respects as well as general prymnesiophyte dominance and low diatom contributions to eukaryotic pigment markers from CHEMTAX analyses (Selph et al., this issue), lower food web structure would seem to be similar between the two spawning regions but rates are up-shifted to higher production and grazing flows in the Argo Basin. Very low diatom production (2-10 mg C m^−2^ d^−1^) was especially noted for the oceanic GoM (Landry et al., 2022a). The large-volume Lugol’s-preserved samples examined by inverted microscopy in the present study gave diatom integrated values of 3.4 ± 1.2 mg C m^−2^ that may involve some cell loss to the acidic preservative but were nonetheless supported by very few diatoms enumerated on EPI slides. For C biomass estimates of this magnitude and FUCO-based growth rates, diatom production in the Argo Basin would be on the order of 1.5 mg C m^−2^ d^−1^ for all cycles, with C3 having the highest value (4.8 mg C m^−2^ d^−1^) from the combination of highest diatom biomass and highest growth rates. These fall in the range of the low diatom production estimates for the GoM.

Process rates in tropical waters that flow from the Indonesian Throughflow into the Argo Basin were previously investigated at 110°E (downstream of the current study location) in late May and early June 2019 (Landry et al., 2022b). For experiments conducted in waters north of 15.5 °S, mean FlChl*a*-based total production (852 ± 149 mg C m^−2^ d^−1^) was slightly higher than our Argo estimate (Table 3), while FCM-based PRO production (430 ± 22 mg C m^−2^ d^−1^) was more than double our February 2022 values, mainly reflecting faster growth rates rather than elevated biomass. With this seasonal difference and with strong winds known to drive upwelling on the northern (Indonesian) side of the Argo Basin during the austral NW Monsoon from June to September, the SBT spawning season likely misses the period of highest regional productivity. Characteristic differences are also evident in comparing the present Argo results to classic open-ocean upwelling regions of the equatorial and eastern tropical Pacific. TChl*a*-based production for the equatorial Pacific (867 ± 96 mg C m^−2^ d^−1^; Landry et al. 2011) was 1.7 fold higher than the mean Argo estimate, and FlChla-based production for the Costa Rica Dome (990 ± 106 mg C m^−2^ d^−1^; Landry et al. 2016) exceeded the Argo estimate by 1.3 fold. More telling, these more productive systems have substantially lower PRO contributions to total productivity (15-20% and ∼6%, respectively, the latter reflecting a picoplankton dominance of *Synechococcus*). Diatom production estimates for the equatorial Pacific (156 ± 21 mg C m^−2^ d^−1^; Landry et al., 2011) were 1.5 to 2 orders of magnitude greater than our crude estimates for diatoms during SBT spawning in Argo Basin.

### 4.3. Microbial food web fluxes during the SBT spawning season

Figure 7 summarizes carbon flux estimates for the microbial food webs in the upper and lower EZ of the southern Argo Basin during SBT larval development based on blended mean values for PRO and the total community with PEUK fluxes determined the Total-PRO difference. The upper EZ incorporates measurements from the surface to 25 m, corresponding to the habitat where SBT larvae reside and feed. The lower EZ incorporates measurements from the lower 3 profile depths, representing a contrasting habitat of microbial food web interactions that can be exploited by mesozooplankton that can move between the lower and upper EZ layers, where they enter the prey fields of larval tuna. The bar heights of all biomasses (mg C m^−3^) and rates (mg C m^−3^ d^−1^) are scaled proportionately to each other and in both upper and lower panels.

**Fig. 7.**
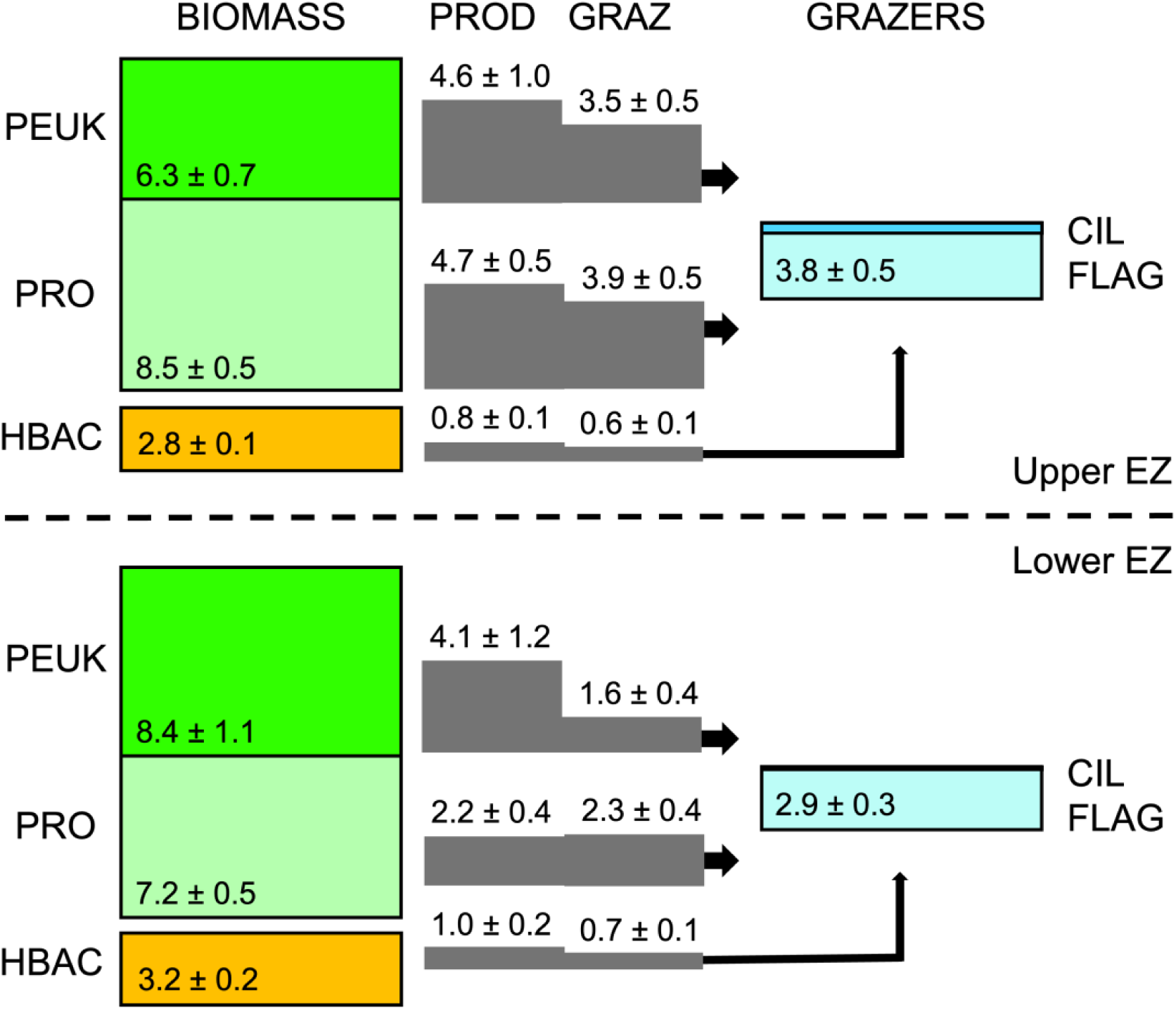
Mean carbon flows within microbial food webs of the upper and lower EZ of the Argo Basin. PEUK = photosynthetic (Chl*a* containing) eukaryotes; PRO = *Prochlorococcus*; HBAC = heterotrophic bacteria. Grazers are ciliates (CIL) and heterotrophic flagellates (FLAG). PROD and GRAZ are C-based rate estimates (mg C m^−3^ d^−1^) for production and grazing, respectively. All biomass (mg C m^−3^) and rate estimates are scaled proportionately by height. Uncertainties are standard errors of mean values (SEM).

PRO is a major contributor to microbial community biomass, production and grazing fluxes in the upper EZ, comprising 48% of the total C of phytoplankton and bacteria, 47% of total C production and 49% of the grazing flux (Fig. 7). The total grazing flux of 8 mg C m^−3^ d^−1^ is 211% of heterotroph C per day, which would give heterotrophic protists an instantaneous growth rate of 0.49 d^−1^ if we assumed a gross growth efficiency of 30% (Straile, 1997). However, at least part of the grazing might be done by mixotrophs within the PEUK (chlorophyll-containing eukaryote) component, which Selph et al. (this issue) found to be abundant in the upper EZ based on FCM cell analyses with a fluorescence tracker of acidic feeding vacuoles and correlated with the abundance distribution of HBAC. Mixotrophic functionality would allow PEUKs to compete with PRO for limiting nutrients in the oligotrophic near-surface waters of the Argo Basin and also contribute to the offset between ^14^C-NCP and Chl*a*-growth based estimates of phytoplankton production. A very rough estimate of the C flow magnitude to mixotrophs might be made by assuming that chlorophyll- and non-chlorophyll-containing protists of similar size should be equally available (sustain similar instantaneous rates of grazing loss) to mesozooplankton in a steady state system. Accordingly, for heterotrophs to experience the same low rate of grazing loss to mesozooplankton as indicted for PEUK (0.16 d^−1^, determined from the difference between PEUK production and mortality loss to protistan grazers), they would require only 27% (2.2 mg C m^−3^ d^−1^) of the total of 8.0 mg C m^−3^ d^−1^ that Fig. 7 suggests is available to them. If the difference in C grazing is assumed to be consumed instead by mixotrophic PEUK, it could account for 38% of the PEUK growth attributed to photosynthesis, assuming that typical 70% losses to egestion, respiration and excretion apply.

On a volume basis, biomasses of PEUK and HBAC increase in the lower EZ relative to surface values, and PRO biomass declines (Fig. 7). Whereas, PEUK and PRO contribute similarly to production in the upper EZ, PEUK production is almost double that of PRO in the lower EZ. HBAC production also increases slightly and in relative magnitude compared to photosynthetic groups. Total grazing of 4.6 mg C m^−3^ d^−1^ corresponds to 159% of heterotroph C d^−1^, which can support an instantaneous growth rate of 0.39 d^−1^ for heterotrophic protists, similar to the 0.40 d^−1^ growth rate estimate for PEUK. Following the same approach for estimating potential phagotrophy by mixotrophs in the upper EZ, protistan heterotrophs require 83% of the total 4.6 mg C m^−3^ d^−1^ grazing fluxes available to them in the lower EZ in order to match the loss rate of PEUK (0.26 d^−1^) to mesozooplankton consumption. As a consequence, mixotroph phagotrophy would support a much lower percentage (6%) of PEUK production in the light-limited lower EZ than in surface waters. This would be consistent with the greater prevalence Chl*a*-containing protists with acidic vacuoles, presumptive of phagotrophic feeding, in the upper versus lower EZ of our study region (Selph et al., this issue).

While major taxa of eukaryotic phytoplankton were not assessed for their individual contributions to C biomass in the present analysis, some inferences can be made about their food web roles from the relative magnitudes of their growth and grazing rates and their relative contributions to carotenoid pigment concentrations. HEX was the dominant carotenoid and had the highest average values of growth and grazing in the upper EZ (Table 2). This leaves little doubt that prymnesiophytes should be major contributors to eukaryotic production and grazing fluxes in this part of the water column, as seen overall in the Gulf of Mexico (Selph et al., 2022; Landry et al. 2022). In the lower EZ, the relative growth ratios of FUCO and BUT compared to HEX were 4-5 times higher than in the upper EZ, and diatom C biomass, though still small, was 4.7 times higher in absolute terms than at shallower depths. These differences would suggest increasing relative roles for diatoms (FUCO) and pelagophytes (BUT) in the deeper stratum.

Lastly, the microbial food web fluxes in Figure 7 put constraints on the resources available for mesozooplankton ingestion when viewed from a steady-state system perspective in which all production is consumed either by micro- or mesozooplankton. At steady state, mesozooplankton consumption is the sum of the net differences between C production and microzooplankton grazing of phytoplankton and HBAC plus the inferred production of heterotrophic grazers. For the upper EZ, this feeding constraint for mesozooplankton is ∼4.5 mg C m^−3^ d^−1^, of which 78% comes from eukaryote production and 33% from Chl*a*-containing prey (nominally the component measured by zooplankton gut fluorescence; Décima et al., this issue). Consideration of C flux through mixotrophic PEUK, as discussed above, would reduce total C flux available for mesozooplankton to 2.8 mg C m^−3^ d^−1^, of which 63% is from eukaryotes and 67% from Chl*a*-containing prey. In the lower EZ, mesozooplankton consumption is constrained to 3.7-4.1 mg C m^−3^ d^−1^, depending on whether flux estimates are taken at face value or interpreted in terms of mixotrophy (the lower value of the range). Of the C available to mesozooplankton in the lower EZ, 89-93% is from eukaryote net production and 61-68% comes from Chl*a*-containing prey.

## 5. Summary and conclusions

Austral summer conditions in the Argo Basin region, downstream of the Indonesian Throughflow and during the peak SBT spawning season, have classic oligotrophic oceanic features with a thick layer of nutrient-depleted surface waters overlying a well-defined DCM. Dilution production rates (527-772 mg C m^−2^ d^−1^) from alternate measurement variables (TChl*a* and FlChl*a*) were consistent overall with lower ^14^C-NCP estimates (447 mg C m^−2^ d^−1^), but offered different interpretations of C cycling efficiency. Interregional comparisons indicated similar structures and relative roles of *Prochlorococcus*, diatoms and prymnesiophytes to those in the bluefin spawning region of the oceanic Gulf of Mexico. Rate estimates were higher than the GoM but substantially lower than tropical open-ocean upwelling regions. PRO was the major contributor to biomass and C fluxes in the upper EZ, where tuna larvae reside. Significant C flows through PEUKs in the upper EZ can accommodate an interpretation in which mixotrophy plays a significant role, aligning with evidence from acid vacuole analyses and community production differences from dilution versus ^14^C-uptake incubations. Process relationships in the lower EZ including the DCM showed a 2-fold C flux decline for PRO, increasing relative contributions from eukaryotes and heterotrophic bacteria, and lower mixotrophy potential. On average, 71% of total production and 86% of PRO production were utilized by consumers in the microbial food web, leaving a residual diet for mesozooplankton substantially richer in eukaryote and heterotrophic (non-Chl*a* containing prey) components than the relative biomass distributions of the microbial community. These results are an initial step to understanding how lower-level food web C flows relate to feeding, growth and development of larval tuna in the Argo Basin.

## Acknowledgements

We gratefully acknowledge captain and crew of R/V *Revelle* and all of the RR2201 science participants for their dedication, professionalism and exceptional contributions to this research during a challenging period of Covid travel restrictions and protocols that greatly extended time away from home and family. This study was supported by U. S. National Science Foundation grants OCE-1851558 (M.R.L.), -1851347 and -2404504 (S.A.K. and M.R.S) and -1851381 (K.E.S.) and is a contribution to the Second International Indian Ocean Expedition (IIOE-2 endorsed project EP046). Seawater and plankton samples were collected under Australian Government permit AU-COM2021-520 and Australian Marine Parks permit PA2021-00062-2 issued by the Director of National Parks, Australia. Views expressed in this publication do not necessarily represent those of the Director of National Parks or the Australian Government. This paper is dedicated to the memory of Prof. Karl Banse (1929-2025): champion of Indian Ocean research and zooplankton grazing processes; influential teacher and generous mentor to generations of young ocean scientists.

## Author statement

M.R.L. conceived the study. M.R.L., M.R.S., N.Y., K.E.S., S.A.K., C.K.F. and R.S. conducted the sampling and experiments at sea. N.Y., K.E.S., S.K. and C.F. provided data from EPI microscopy, flow cytometry, ^14^C-NCP and fluorometric Chl*a*, respectively. R.B. and R.S. contributed biomass analyses from Lugol’s samples. M.R.L. analyzed results and wrote the manuscript. All authors contributed to comments and edits of the manuscript.

